# QTL identification and characterization of the recombination landscape of the mountain pine beetle (*Dendroctonus ponderosae*)

**DOI:** 10.1101/2025.04.08.647844

**Authors:** Camille Pushman, Gregory J. Ragland, Michael E. Pfrender, Barbara J. Bentz, Ryan R. Bracewell

## Abstract

Insect pests can rapidly accumulate in number and thrive in diverse environments, making them valuable models for studying phenotypic plasticity and the genetic basis of local adaptation. The mountain pine beetle (*Dendroctonus ponderosae)* is a major forest pest and adult body size and generation time are two traits that vary among populations and directly influence reproductive success and outbreak dynamics. To identify regions of the genome linked to these two traits, we generated ddRAD sequencing data from an F_2_ intercross, using populations from two Y haplogroups with phenotypic and genetic differences in these traits. A high-density linkage map was generated and QTL analyses performed. We identified a single large effect QTL for generation time, associated with an adult diapause. The QTL spans the entire X chromosome, peaking over the evolutionarily conserved portion of the X. We were unable to detect a significant QTL for body size. Our linkage map identified putative inversions shared by parents that are absent in the published reference genome, with three putative inversions on chromosomes 2, 3, and the X. We also detected extensive regions of low recombination that were associated with low gene density, indicative of large pericentromeric regions. Surprisingly, we found that in our cross, F_2_ males inherited X chromosomes with significantly fewer crossover events than F_2_ females. Our findings provide information about the recombination landscape, the sex-biased inheritance of recombined X’s, and the genomic location of a key trait in a major forest pest.

## INTRODUCTION

Insect pests that go through periodic population outbreaks are valuable models for studying adaptation and the underlying genetic changes that facilitate their abundance (Zhu-Salzman and Zeng 2008; Pélissié et al. 2018; McCulloch and Waters 2023). Insects prone to outbreak can typically thrive in a diverse range of environmental conditions, offering insights into how species quickly respond through phenotypic plasticity and can adapt over time (Petegem et al. 2016; Doellman et al. 2020; Dowle et al. 2020). Exploring the genetic basis of such complex traits can be achieved through quantitative trait loci (QTL) analysis, which uses genotype data from recombinant individuals to create genetic linkage maps and identify regions of a genome associated with a specific phenotype. QTL approaches have successfully identified loci associated with a variety of traits in insects, including photoperiodism (Bradshaw et al. 2012), cold tolerance (Königer et al. 2019), and wing patterning (Jones et al. 2012). QTL analyses are most informative when using high-density linkage maps (Dhingani et al. 2015) and restriction-site associated DNA sequencing (RAD-seq) provides a cost-effective method for identifying numerous genetic markers in non-model systems (Davey et al. 2011; Andrews et al. 2016).

In addition to their application in QTL analyses, linkage maps are a useful tool for examining patterns of recombination across the genome (i.e., the recombination landscape). Recombination influences genetic linkage between alleles (Yeaman 2013), genetic diversity (Wright et al. 2006), and mutational load (Jay et al. 2022) Recombination rates are also thought to be under strong selective pressure (Ortiz-Barrientos et al. 2016). Recombination patterns and rates have been found to vary between species, populations, and sexes (Kong et al. 2010; Wang et al. 2023). A variety of mechanisms can modify patterns of recombination such as chromosomal inversions (Hoffmann and Rieseberg 2008), transposable elements (Rizzon et al. 2002; Kent et al. 2017), chromatin organization (Jin et al. 2021), and centromeres (Copenhaver et al. 1999; Myers et al. 2005). Chromosomal inversions are particularly interesting in the context of adaptation because they have been found to facilitate trait divergence, as seen for body size and development time in *Drosophila* (Betrán et al. 1998), mate preference in *Anopheles* (Ayala et al. 2013), and flowering time in *Mimulus* (Lowry and Willis 2010).

The mountain pine beetle (*Dendroctonus ponderosae* Hopkins, MPB) is a native bark beetle in western North America. MPB has eruptive population dynamics (Raffa et al. 2008) and an expansive distribution extending from northern Baja California to northern British Columbia and Alberta, Canada (Dowle et al. 2017). Among MPB populations, there is plasticity and genetic variation in thermally regulated traits that have direct links to rapid demographic change. This variation makes MPB an excellent model for exploring the genetic architecture of adaptation (Bentz et al. 2011; Janes et al. 2014; Dowle et al. 2017). MPB attack and reproduce in most species of *Pinus* and tree death is typically required for brood success. Highly defended and food-rich trees are only accessed using synchronous adult attacks (Boone et al. 2011).

Synchronous attacks, in turn, are dependent upon synchronous and seasonally appropriate brood adult emergence from previously attacked trees, and generation timing is critical (Logan and Bentz 1999). Populations from the southern part of the MPB distribution, including Arizona (AZ), were found to have longer generation time than more northern populations in common gardens, suggesting adaptation to local climate (Bentz et al. 2001; Bracewell et al. 2013). In a field-based reciprocal translocation experiment, Soderberg et al. (2021) confirmed a longer generation time in an AZ population compared to a population from northern Utah (UT), highlighting the phenotypic and genetic variation between the populations. Adult body size, which is positively correlated with multiple traits known to influence MPB survival (McGhehey 1971; Safranyik 1976; Elkin and Reid 2005; Bentz et al. 2011), was also shown in the same common gardens to vary among populations. Adult body size and generation time are heritable traits that are likely under strong selection given their variability among populations and importance to rapid population growth (Bentz et al. 2001).

In addition to population-level variation in life history traits, MPB also has distinct neo-sex chromosomes that show substantial genetic variation across the range of the species (Bracewell et al. 2017). Neo-sex chromosomes occur when an ancestral sex chromosome fuses to an autosome (Zhou and Bachtrog 2012). The neo-sex chromosomes of MPB appear rather large in chromosome squashes (Lanier and Wood 1968) and the neo-X has been found to make up at least 20% of draft genome assemblies (Keeling et al. 2022; Bracewell et al. 2024). After a fusion, neo-sex chromosomes can remain similar at the DNA sequence level, and only begin to differentiate when recombination stops between the neo-X and neo-Y (Zhou and Bachtrog 2012; Zhou et al. 2012; Ponnikas et al. 2022; Akagi et al. 2023). In the case of MPB, the spatial pattern and extent of neo-X/neo-Y differentiation is currently unclear and there is no estimate of when these chromosomes may have formed. Additionally, crosses between populations with distinct neo-Ys produce sterile males, suggesting these chromosomes could be important in speciation (Bracewell et al. 2011; Bracewell et al. 2017).

The current MPB distribution is made up of three distinct neo-Y haplogroups (termed ‘eastern’, ‘western’ and ‘central’), that are geographically restricted and associated with reproductive incompatibilities and population genetic structure (Bracewell et al., 2017; Dowle et al., 2017). Previous studies have shown crosses between western x eastern and western x central populations produce males with highly reduced sperm quantities (Bracewell et al. 2011; Bracewell et al. 2017). The northern UT and AZ populations that were previously found to have phenotypic and genetic differences in generation time and body size (Table 1) reside in the eastern and central haplogroups, respectively (Dowle et al., 2017). Our overall goal was to examine potential reproductive incompatibilities between the AZ and UT populations and to further examine the underlying genetic basis of trait differences between the eastern and central haplogroups using QTL analysis. To achieve this, we first characterized chromosome-wide patterns of sequence similarity between the neo-X and neo-Y. This analysis provides insight into the structure and differentiation of these chromosomes and serves as a foundation for accurate genotyping and QTL mapping. Next, we applied double-digest RAD (ddRAD) sequencing to create a high-density linkage map and identify QTLs associated with body size and generation time. Finally, we leveraged the recently published chromosome-level MPB genome assembly (Keeling et al. 2022) to compare genetic and physical distances using Marey maps (Chakravarti 1991). These comparisons allowed us to detect putative inversions, identify pericentromeric regions, and examine sex-biased inheritance of non-recombining regions of the X chromosome.

**Table 1:**
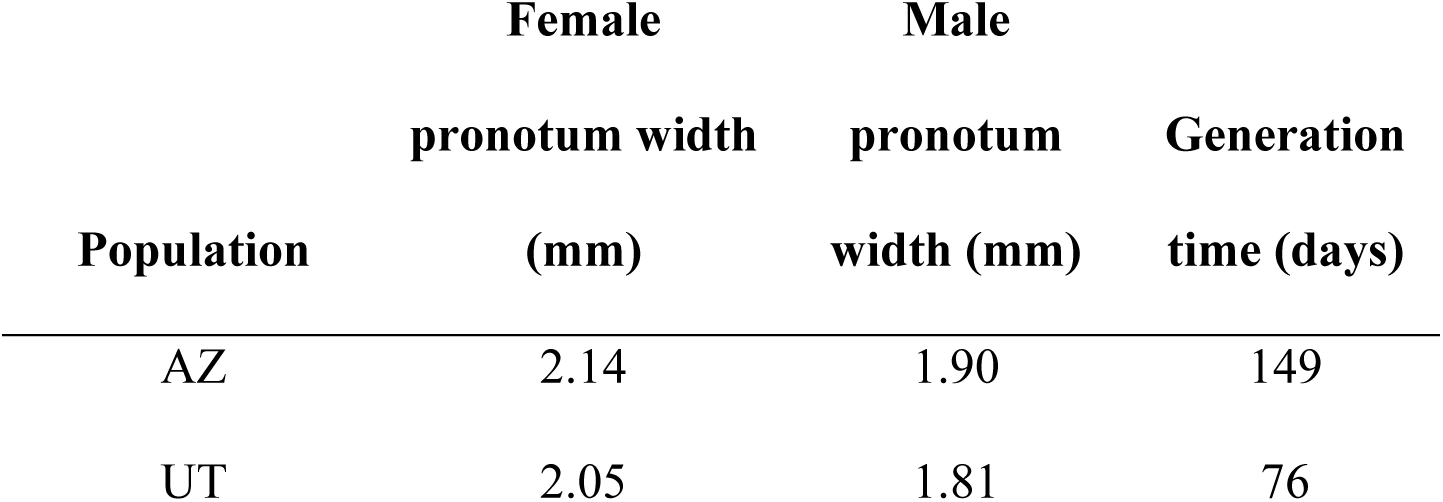
Estimates of mean body size (pronotum width) and generation time of AZ and UT populations from Bracewell et al. (2013). Our AZ and UT populations were sourced from locations near Bracewell et al. (2013).

## METHODS

### MPB collections and crosses

We collected beetles from lodgepole pine (*Pinus contorta)* in the Wasatch-Cache National Forest, UT (41° 58’ N, -111° 31’ W), and southwestern white pine (*P. strobiformis*) in the Coronado National Forest, AZ (32° 38’ N, -109° 52’ W). Infested trees were harvested geographically near populations used in previous studies showing genetic differences in adult body size and generation time (Table 1) (Bracewell et al. 2013; Soderberg et al. 2021). Trees infested with beetles were felled, cut into 30 - 45cm sections, and sealed with paraffin wax. The sealed tree sections (hereafter referred to as bolts) were placed in rearing containers and stored at room temperature (∼22 °C) until beetles emerged. Brood adults were moved to Petri dishes lined with moist filter paper and stored at ∼3 °C for up to 20 days.

We performed crosses in uninfested lodgepole pine bolts from the Wasatch-Cache National Forest, UT. For each cross, holes were drilled into the phloem layer of the bolt, so the mating pair could be inserted, and then covered with fabric mesh to prevent escape. We placed the bolts in rearing containers at room temperature (∼22 °C), on a ∼ 9-hour light:15-hour dark photo regimen. Adults used in all crosses were randomly selected from the peak emergence period (∼15 days of highest cumulative number of beetles).

Our P_0_ cross was between an AZ male and UT female. We collected the resulting F_1_ offspring every two days and stored them using methods described above. Thirty F_1_ offspring were randomly selected and paired to complete 15 intercrosses. We recorded sex, body size, and day of emergence for all F_2_ offspring. Beetles were sexed using established protocols (Lyon 1958). Body size was estimated by measuring pronotum width, a common proxy for overall body size in bark beetles (McGhehey 1971; Amman 1982). Generation time was estimated as the number of days from when the mating pair was first inserted into the bolt to when the brood adult emerged.

### DNA extractions and sequencing

To extract DNA from P_0_ parents and all 243 F_2_ individuals we used thorax tissue and a Gentra Puregene tissue kit (Qiagen, Velencia, CA). Extracted DNA (90-110 ng) was transferred to a digestion reaction using EcoRI and Msel restriction enzymes (New England Biolabs, NEB), adapters were ligated to DNA fragments, and PCR-amplified for 30 cycles using adapter- complimentary primers. We visualized PCR products on an agarose gel to verify proper amplification. We selected 300-500 bp fragments using pooled DNA (∼96 individuals) and a BluePippin System (Sage Science, Beverly, MA), then cleaned them using Ampure beads (Ampure beads Beckman Coulter, Indianapolis, IN). Libraries were assessed using a Bioanalyzer 2100 system (Agilent) and sent for sequencing at BGI Americas Inc. in Davis, CA. An Illumina HiSeq 2000 (v3 chemistry) was used to complete 100bp paired-end sequencing across three lanes. Our ddRAD sequencing data was generated alongside data described in (Dowle et al. 2017).

### Data filtering

The 100bp paired-end reverse reads were of poor quality so we followed methods described in Dowle et al. (2017), that suffered the same issue and therefore used only the forward reads in our analysis. Preliminary FastQC results indicated forward reads were of high quality and adequate for map construction and QTL analysis. All forward reads were processed and demultiplexed with Stacks version 1.48 (Catchen et al. 2013) *process-radtags* using default settings and a minimum sequence length of 10bp. During this step, we also removed reads without a base call or with a low-quality score. The resulting files were mapped to the female MPB reference genome (Keeling et al. 2022; Accession GCA_020466585.2) using BWA-MEM (Li 2013) with split reads marked as secondary. Mapped reads were sorted with SAMtools (default parameters)(Danecek et al. 2021), formatted using Picard AddOrReplaceReadGroups using the lenient validation stringency option (Picard toolkit 2019), and indexed using SAMtools.

Given the large neo-sex chromosomes of MPB (Bracewell et al. 2024), we first explored neo-X/neo-Y differentiation using whole genome sequencing (WGS) data to determine how to appropriately handle genotyping (below) since neo-Y reads could map to the X chromosome. We compared previously published Illumina whole genome sequencing data derived from a single male and female (samples SRS2676974, SRS2676969) and mapped the reads to the female MPB reference genome (Accession PRJNA638278) with BWA-MEM (Li 2013). We used SAMtools (Danecek et al. 2021) to filter (mapping quality > 30), sort, and index the sequences and BEDtools (Quinlan and Hall 2010) and a custom Perl script to estimate average X chromosome coverage in 25 kb windows. These results were graphically visualized (Wickham and Wickham 2016).

We called single nucleotide polymorphisms (SNPs) using GATK version 3.8 UnifiedGenotyper (McKenna et al. 2010) and filtered them using VCFtools version 0.1.13 (genotype quality > 30, depth > 20, biallelic sites, remove insertions/deletions) (Danecek et al. 2011). We identified sites where P_0_ individuals were homozygous for different alleles, and analyzed these sites in F_2_ offspring using VCFtools (Danecek et al. 2011). This also allowed us to determine what alleles came from each parent, which we will refer to as AZ and UT haplotypes hereafter, for the AZ and UT parents. Only sites that mapped to the largest 12 chromosome-level scaffolds in the reference genome were retained. We also removed markers that were missing genotypes for ≥ 68% of individuals, and individuals missing data at ≥ 99% of markers. These cutoffs were chosen after exploring a range of thresholds and allowed us to maximize the number of individuals in the mapping panel without significantly compromising marker number and map quality. To reduce redundant markers we used VCFtools to thin the dataset and retained 1 marker per 100 kb.

After converting our VCF to R/qtl format and importing, R/qtl detected 1,042 genotype calls on the X chromosome that should not occur based on our cross design (estimate.map = FALSE, convertXdata =TRUE) (Broman et al. 2003). These were all males with heterozygous genotype calls or homozygous females with two AZ-type X chromosomes. Because the F_1_ male had to transmit one, non-recombined, UT-type X chromosome to female offspring, F_2_ females should not have two AZ-type X chromosomes. Similarly, the F_2_ males cannot have a heterozygous genotype because they are hemizygous for the X chromosome when the Y is degenerated. We found that all these flagged genotypes fell into two broad categories: low-quality markers or mapping error. Any marker where multiple individuals had a flagged genotype at that single marker (which would require a double crossover event) was considered low quality and omitted. The remaining flagged genotypes localized to the first ∼26 Mb of the X chromosome and were only found in males, which is same region where the neo-X and neo-Y coverage appear to be most similar (see RESULTS). Thus, these genotypes likely resulted from the neo-Y mapping to the neo-X chromosome, and we used the *convertXdata* parameter to remove the flagged genotypes. In a previous study, a similar pattern of genotyping errors over this region of the X was also resolved using similar methods (Keeling et al. 2022).

### Linkage map construction

Low quality markers complicate linkage mapping and we took several steps to remove them. First, we filtered out any markers flagged as possible genotyping errors (error.prob = 0.001). Markers that displayed significant segregation distortion (R/qtl geno.table function with cutoff of *p* < 0.01) were also excluded, except on the X chromosome. We did not exclude markers on the X chromosome during this step because male hemizygosity on the X would appear as homozygous genotype calls, which skew the genotype distributions away from Mendelian expectations. Marker order was verified using the R/qtl orderMarkers function (without rippling). As a final validation, we used heat map associations to identify markers in suboptimal locations and manually moved them to their location with highest linkage association (per R/qtl recommendations). Specifically, we altered the ordering of markers when linkage suggested an improved location. Any markers that showed clear linkage across more than one chromosome via heat map associations were removed. Our linkage map was constructed by using the est.map function in R/qtl with default parameters.

In addition to our combined linkage map which included F_2_ male/female data, we constructed sex-specific linkage maps to explore patterns of recombination along the X chromosome. We did this by repeating the final est.map step (as described above) separately for F_2_ males and F_2_ females. Our sex-specific linkage mapping required no normalization steps since both sexes inherit only one recombined X chromosome from their F_1_ mother. As a result of the crossing design, the naïve expectation is that both sexes should show similar recombination patterns across all 11 autosomes and X chromosome. In addition to comparing our maps, we completed Wilcox-rank sum tests with Bonferroni corrections to determine whether F_2_ males and females differed significantly in the number of estimated crossover events on any chromosome.

### QTL analysis and candidate gene identification

We performed a single QTL analysis of generation time and body size using standard interval mapping in R/qtl (n.perm = 1000, perm.Xsp = TRUE, alpha = 0.05, method = “em”). Logarithm of odds significance cutoffs were determined using the scanone output (perm.Xsp = TRUE, alpha = 0.05). We used an approximate Bayesian credible interval with a probability cutoff of 0.95 to define the peak of the QTL and located the corresponding physical locations of the peak markers using the reference genome coordinates. To explore whether particular gene ontologies were overrepresented in our peak QTL region, we performed a gene ontology (GO) enrichment analysis in g:Profiler using any protein-coding genes that overlap with the QTL region according to the MPB reference annotation (Accession GCA_020466585.2) with a significance cutoff of *p* < 0.05.

### Visualizing the recombination landscape using Marey maps

Marey maps were created by plotting physical position (genome base-pair) against genetic position (centimorgan, cM) for each marker. We defined putative inversions as >10 consecutive markers where physical location (bp) decreased as the genetic map (cM) increased. The endpoints were assigned using the maximum and minimum physical location of the markers within the putative inversion. In our study, a putative inversion could represent a real segregating inversion where both the UT and AZ P_0_ share a unique chromosomal configuration not found in the published reference genome, or in contrast, could represent a potential scaffolding mistake in the reference genome. Indeed, previous work scaffolding the MPB genome was challenging with some placements having mixed support (Keeling et al. 2022). To better understand this potential population-level variation in the recombination landscape, we also visualized linkage map data from Keeling et al., (2022), which was used to scaffold the reference genome, and was constructed from crosses using MPB collected in Canada. To search for any transmission distortion, we created and compared Marey maps between F_2_ males and females. Gene density across chromosomes was determined as the number of protein coding genes in 500 kb windows (Accession GCA_020466585.2). The local recombination rate at each marker was estimated using a local regression (span 0.7) via the MareyMap R package (Rezvoy et al. 2007). Because putative inversions (as described above) would cause a false negative recombination rate for multiple markers over large physical distances, we accounted for this by reversing marker order within putative inversions. Recombination rates of each marker were plotted against gene density, and we applied a linear regression to test the significance of this relationship. Seven markers, which did not follow the pattern of surrounding markers and skewed recombination rates to be negative, were excluded. Despite our filtering efforts, some markers maintained a negative recombination rate when compared to adjacent markers, a pattern not uncommon in genetic maps (e.g., (Akopyan et al. 2022; Keeling et al. 2022). We retained these markers to avoid over-filtering the dataset.

## RESULTS

### MPB X chromosome and linkage map

Consistent with MPB having ‘young’ neo-sex chromosomes, we found that male neo-Y reads readily mapped to the neo-X chromosome when using a female genome assembly (Figure 1). However, the extent of mapping varied with respect to chromosome location suggesting varying levels of neo-X/Y differentiation (Figure 1). The region from 48 Mb to the end of the chromosome at 64 Mb had roughly double the sequencing coverage in females when compared to males, as expected for the hemizygous ancestral-X portion of the X chromosome (Figure 1). The middle portion of the X chromosome (∼26 - 48 Mb) showed intermediate male/female differences, while read coverage was somewhat similar over the first ∼26 Mb of the X. The observed sequence similarity of the first ∼26 Mb of the neo-X helped guide our filtering (see METHODS).

**Fig. 1.**
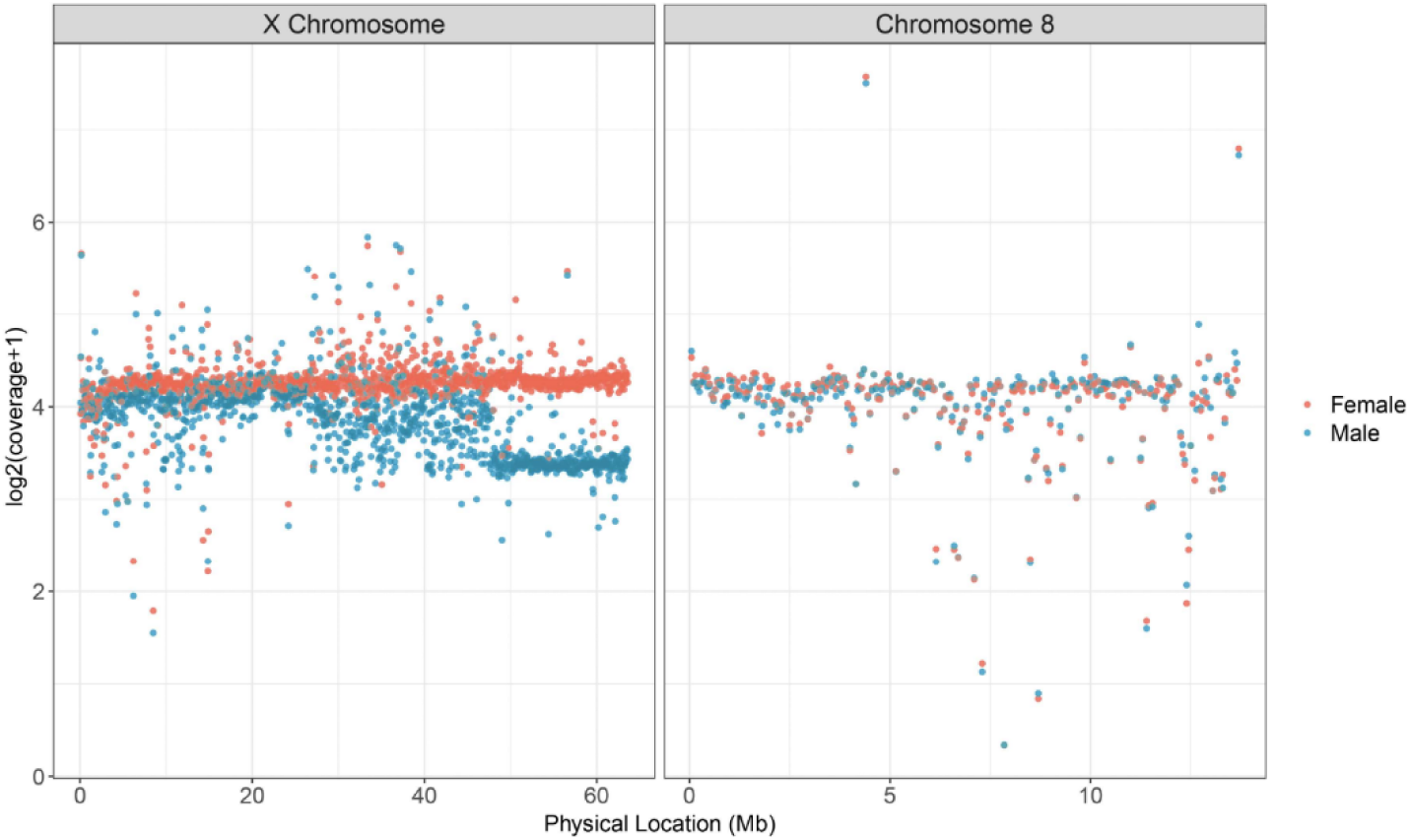
Male (blue) and female (red) Illumina whole genome sequencing data mapped to the X chromosome and a representative autosome (chromosome 8). Each point represents normalized average coverage over a given 50 kb window.

We found no reproductive incompatibility between the populations, and the P_0_ cross (UT female x AZ male) produced viable and fertile F_1_ individuals. Of these, 15 male/female pairs were used to produce 243 F_2_ offspring (132 males and 111 females). Sequencing of 245 samples (all F_2_ individuals and both P_0_ individuals) returned an average of 1.13 ± 0.705 million reads per individual. Of these, an average of 1.08 ± 0.667 million reads (95%) per sample mapped to the reference genome. After filtering, P_0_ individuals had a homozygous genotype call of alternate alleles at 10,209 SNPs. We called these SNPs in the F_2_ offspring, leaving us with 10,069 informative sites. We removed 90 SNPs that did not align to the largest 12 scaffolds in the reference, 97 low-quality SNPs (< 75 genotypes), 62 low coverage individuals (< 100 total genotype calls) and thinned the dataset (kept 1 site per 100,000 bp), leaving 822 SNPs from 181 individuals (101 male, 80 female) for our final dataset. Our final linkage map consisted of 12 linkage groups, corresponding to 11 autosomes and one X chromosome (Figure 2). The total map length was 1069.2 cM, with an average linkage group size of 89.1 cM.

**Fig. 2.**
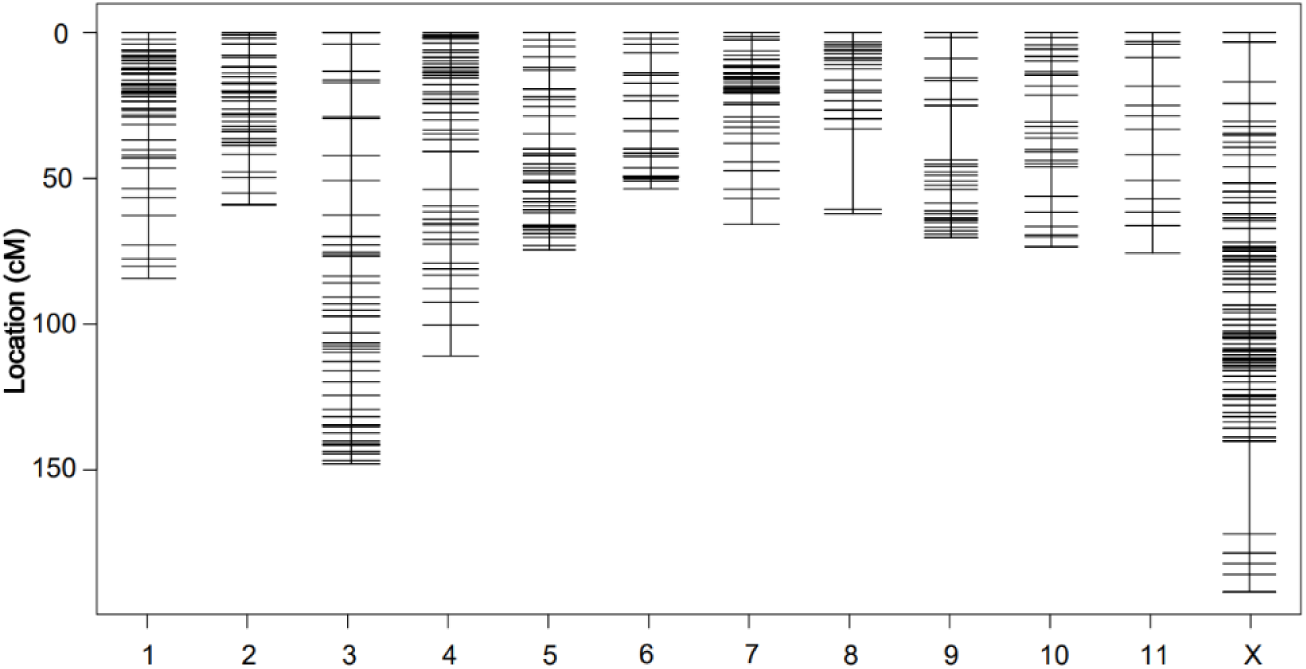
Linkage map consisting of 11 autosomal linkage groups and an X linkage group. The Y axis shows marker location in centiMorgans.

### QTL analysis, and candidate gene identification

We identified one significant QTL for generation time (LOD = 29.8, *p* = < 0.0001, Figure 3A). Markers above the significance threshold occur across 61.8 Mb of the X chromosome (1.5 to 62.3 Mb), spanning the neo-X and ancestral-X portions of the X chromosome (Figure 3A). This significant region includes 3,388 annotated protein coding genes. To narrow the scope of our analysis to a more manageable size, we focused our analysis on the 95% confidence interval (54.9 Mb to 59.7 Mb) which contains 211 annotated protein coding genes. This region was not found to be significantly enriched for any annotated GO terms. A long generation time (> 120 days) was only detected in males hemizygous for the AZ-haplotype allele at the peak marker (Figure 3B). No homozygous AZ-haplotype females were produced due to our crossing design. Heterozygous females displayed generation times more similar to UT-haplotype (Figure 3B), suggesting some dominance of the UT-haplotype short generation time. No significant QTL was detected for body size.

**Fig. 3.**
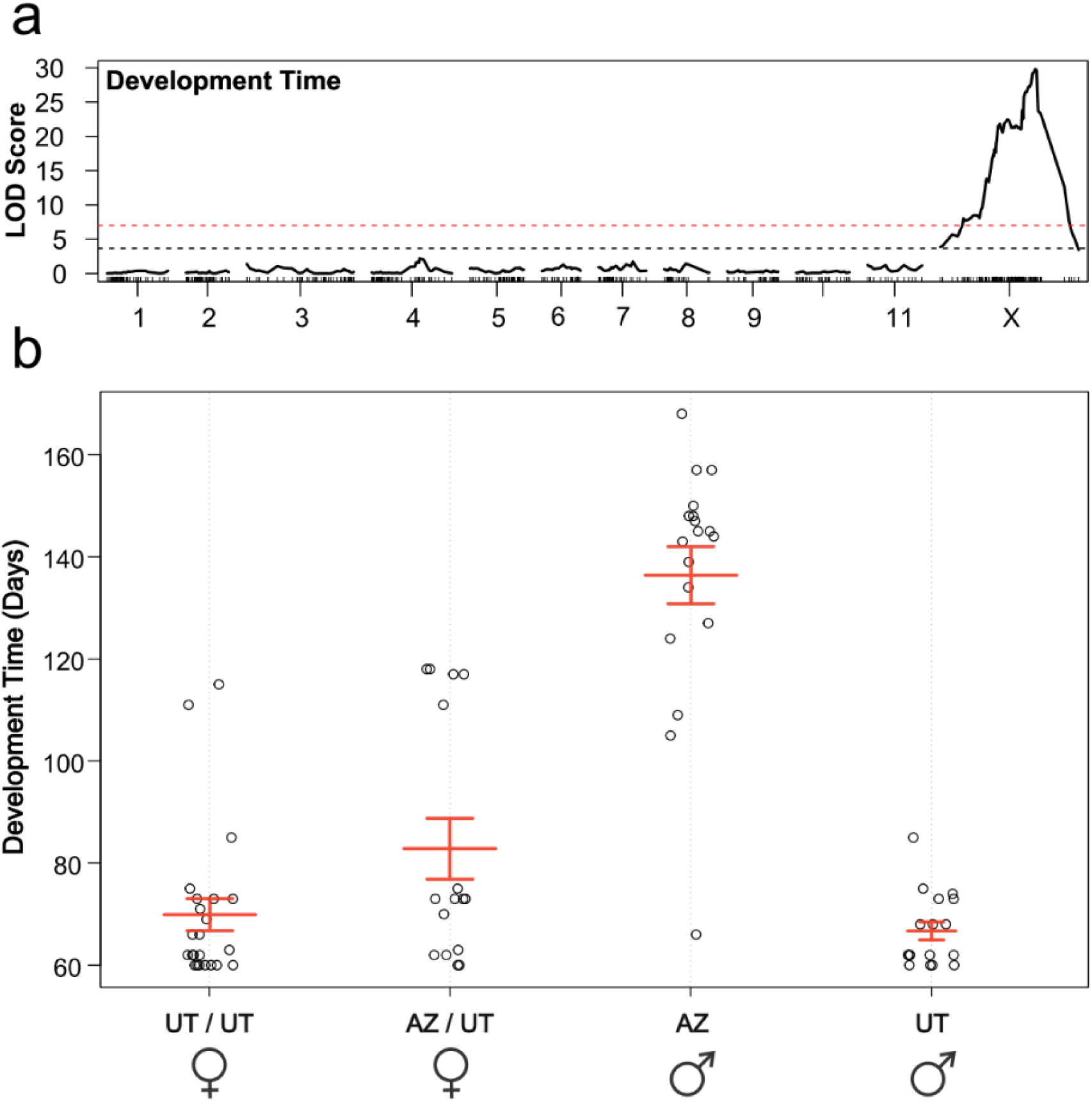
a) QTL analysis for generation time. Logarithm of odds (LOD) scores for markers shown (black solid line) including significance cutoffs for the autosomes (dashed black, LOD=3.67) and X chromosome (dashed red, LOD=7.02). b) Generation time of F_2_ individuals by genotype and sex at the peak QTL marker (58.3 Mb on X chromosome). Whisker boxes represent the interquartile range.

### The recombination landscape

Marey maps showed a complex recombination landscape across MPB chromosomes (Figure 4). First, relative to the published reference genome, we found putative inversions on chromosome 2 (3.7 Mb long), chromosome 3 (5.5 Mb long), and the neo-X chromosome (18.0 Mb long) (Figure 4). The identification of a putative X inversion was initially intriguing given the overlap with the development time QTL however we believe this is likely a reference genome scaffolding issue (discussed below). Second, we uncovered large regions (> 10 Mb) of low recombination on chromosomes 1, 4, 5, 6, and 7 (Figure 4). When comparing all chromosomes, we found a significant relationship where regions of low recombination were correlated with regions of low gene density (R^2^ = 0.295, *p* < 0.0001; Figure 5, Supplemental Fig 1). Regions of low recombination, and low gene density, tended to occur at the ends of all chromosomes (Figure 4), except the X, suggesting all autosomes are likely acro or telocentric and the X is likely metacentric. The low levels or recombination seen across the middle portion of the X in both our data and Keeling et al. (Figure 4) likely contribute to the broad development time QTL, and possible mis-scaffolding.

**Fig. 4.**
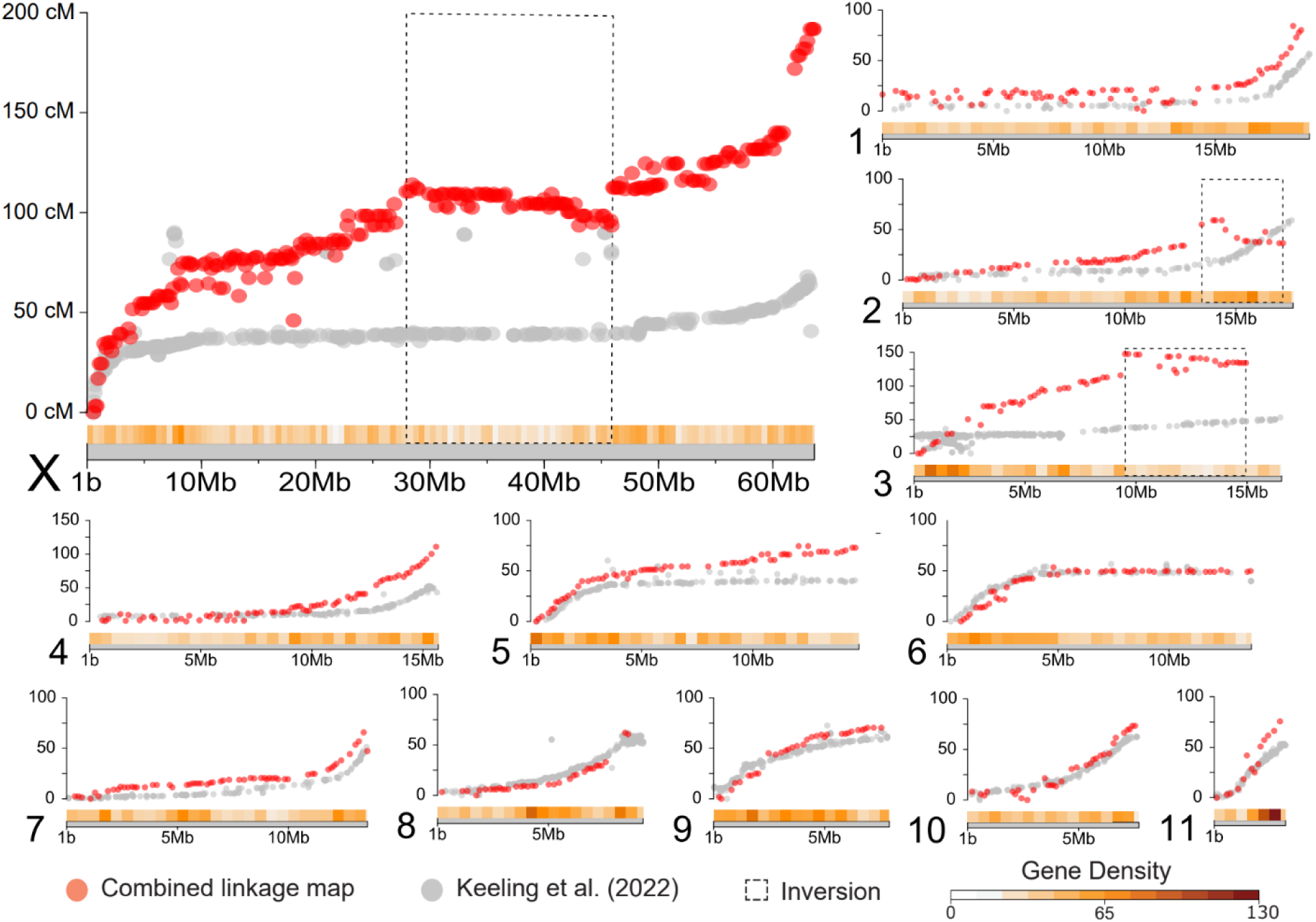
Marey maps of our combined data (F_2_ male + F_2_ female; red) and data from Keeling et al. 2022 (grey). Inversions are shown in dashed boxes. Each point represents a comparison of the marker’s physical position on the X axis (in megabases) and genetic position on the Y axis (in centiMorgans).

**Fig. 5.**
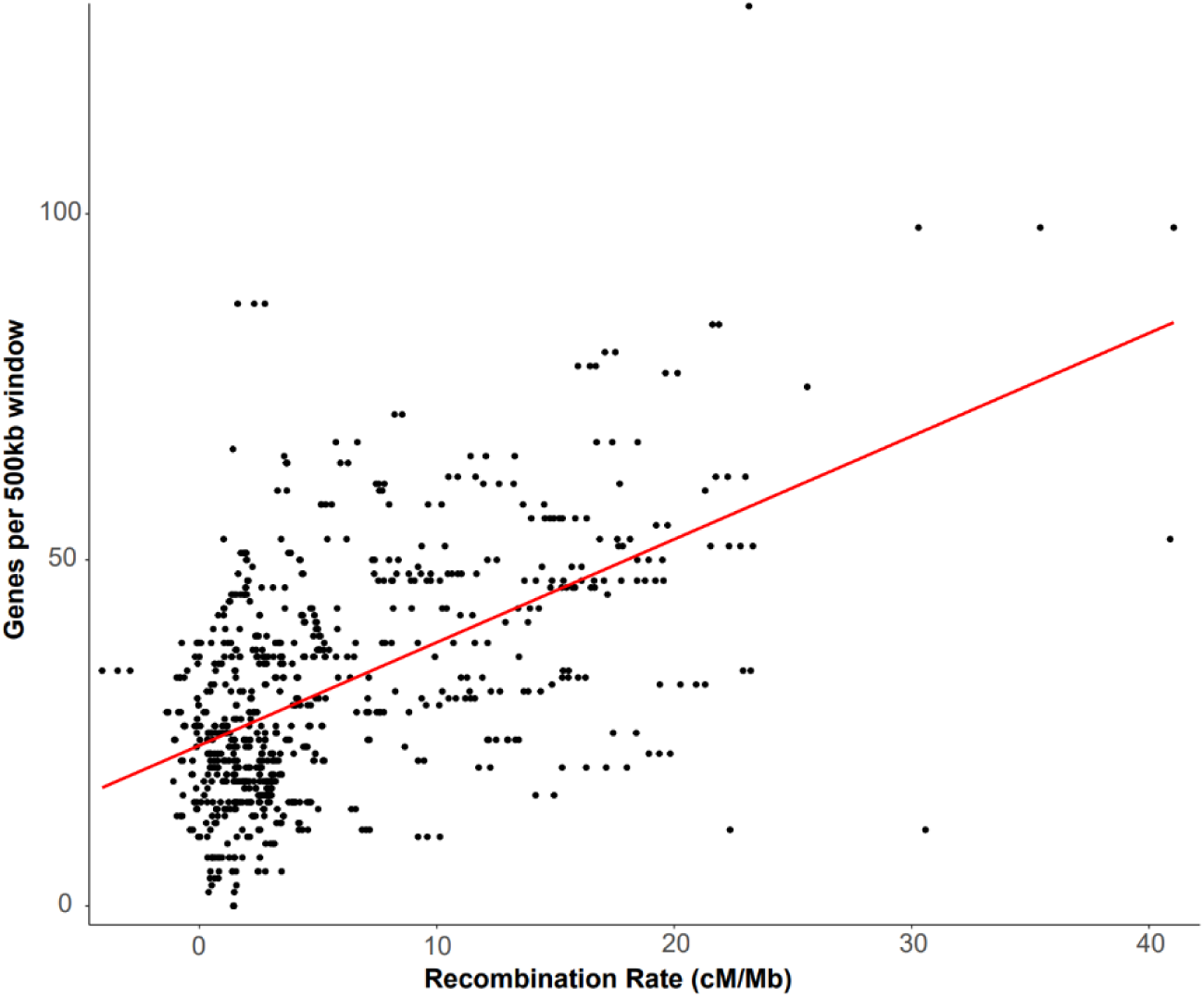
Linear regression of recombination rates (centiMorgans per megabase) against gene density (genes per 500 kb window). Each point represents a marker in our final linkage map. Red line is the fitted linear regression.

Our female-specific and male-specific linkage maps, which should be roughly equivalent based on our methods, measured 1166.3 cM and 1019.1 cM, respectively. The 142 cM difference was driven by the X chromosome, which was 118.3 cM longer in the female map (Figure 6A).

**Fig. 6.**
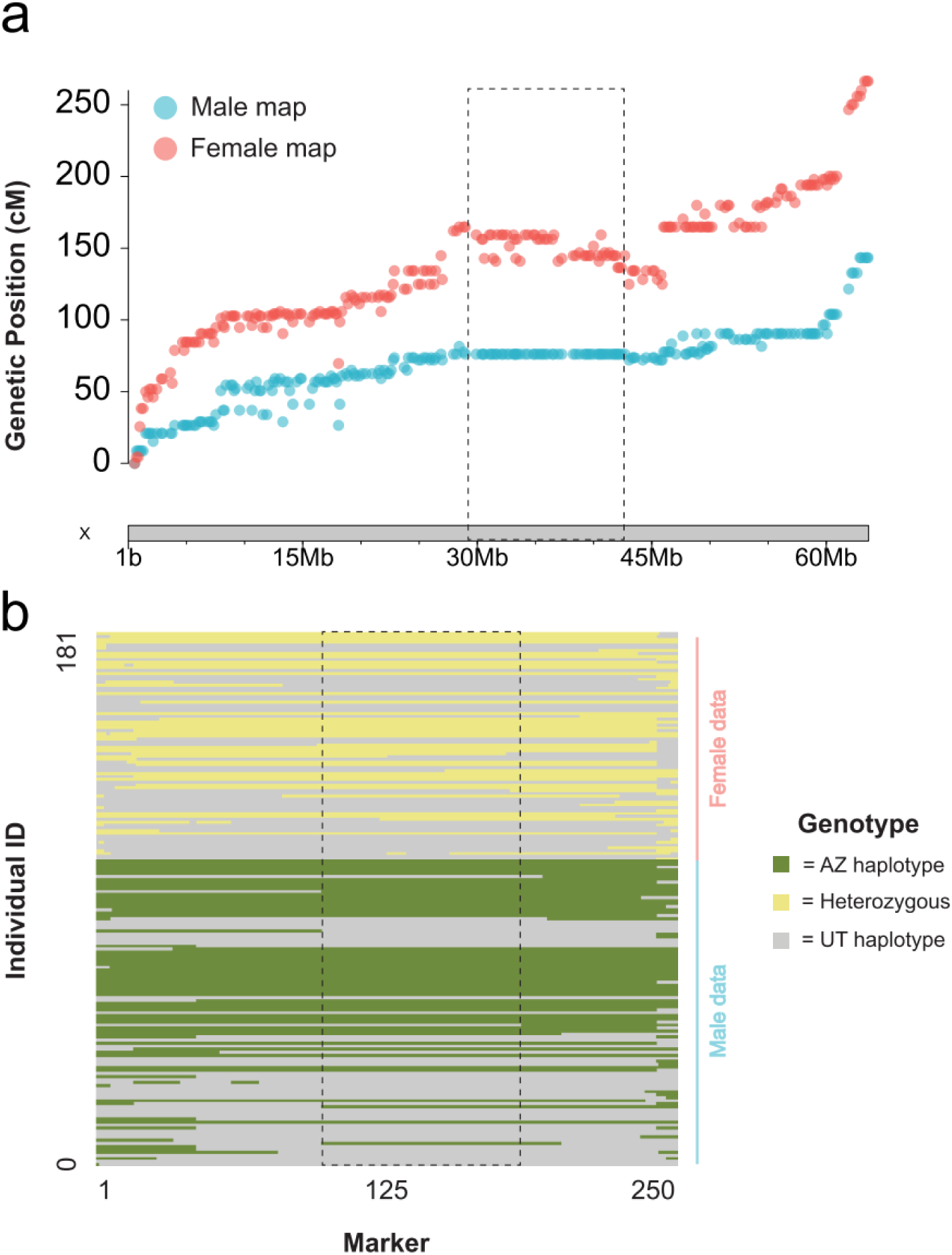
a) Marey map for the X linkage group, comparing F_2_ female (red) and F_2_ male (blue) linkage maps. The region where male X chromosomes show no evidence of a crossover are highlighted with a black dashed line. b) Inferred haplotypes of all F_2_ individuals across the X chromosome. Each row represents an individual, and each column represents a marker in the linkage map for the X chromosome. Dashed box highlights same region as in a).

We found that F_2_ females had significantly more crossover events than those found in F_2_ males (Wilcoxon rank-sum *p* < 0.001, W = 5448, Table 2). Across the X chromosome, females inherited a chromosome with an average of 1.8 crossover events while males inherited an average of 0.8 (Table 2). No autosomes showed a significant difference in recombination events between the sexes (Table 2). To explore this further, we repeated these analyses by first excluding the first 25 Mb of the X chromosome since it required special filtering (See Methods) (Wilcoxon rank-sum *p* < 0.001, W = 5497), and second by analyzing only the ancestral-X portion of the chromosome since genotyping cannot be influenced by the neo-Y (Wilcoxon rank-sum *p* < 0.001, W = 5223). Both analyses found significantly more recombination events on the X chromosome found in F_2_ females. Inspection of haplotype patterns show all F_2_ males lacked a crossover in the region from 26.9 – 49.0 Mb (n = 101), while 8% of F_2_ females had at least one crossover (n = 6 of 80) (Figure 6B). Interestingly, although there was a lack of recombination in this region, the haplotypes segregate evenly, and we observed a near-equal frequency of the AZ (n = 54) and UT (n = 47) haplotypes in F_2_ males (Figure 6B).

**Table 2:** Mean and standard error of crossover events found in F_2_ males and females and results of Wilcoxon-rank sum test corrected for multiple comparisons. NS = not significant.

| Chromosome | Adjusted p-value | Mean XO Events<br>(Females) | Mean XO Events<br>(Males) |
| --- | --- | --- | --- |
| X | 0.0002 *** | 1.8 ± 0.2 | 0.79 ± 0.1 |
| 1 | NS | 1.05 ± 0.1 | 0.91 ± 0.1 |
| 2 | NS | 0.96 ± 0.1 | 0.78 ± 0.1 |
| 3 | NS | 1.9 ± 0.2 | 1.6 ± 0.2 |
| 4 | NS | 1.4 ± 0.2 | 1.2 ± 0.2 |
| 5 | NS | 1.2 ± 0.1 | 0.95 ± 0.1 |
| 6 | NS | 0.55 ± 0.1 | 0.56 ± 0.1 |
| 7 | NS | 0.89 ± 0.1 | 0.77 ± 0.1 |
| 8 | NS | 0.60 ± 0.1 | 0.58 ± 0.1 |
| 9 | NS | 0.80 ± 0.1 | 0.76 ± 0.1 |
| 10 | NS | 0.84 ± 0.1 | 0.79 ± 0.1 |
| 11 | NS | 0.62 ± 0.1 | 0.84 ± 0.1 |

## DISCUSSION

In this study, we identified a significant QTL for generation time that is associated with an adult diapause phenotype in the AZ haplotype (central MPB haplogroup). We also characterized the recombination landscape across the genome and examined spatial patterns of neo-X/neo-Y differentiation in MPB. We identified regions of low recombination and low gene density on multiple autosomes, which are indicative of pericentromeric regions. Additionally, we compared recombination patterns between F_2_ males and females. This comparison showed a reduced recombination rate on the X chromosome in all males and inheritance of a crossover-free 15 Mb region of the neo-X. Our results lay the groundwork for future studies of the evolution of diapause, the role of structural variation in adaptation, and provide much needed information about beetle genome structure and patterns of recombination.

The QTL peak associated with generation time on the X chromosome spanned nearly the entire chromosome, making it difficult to identify putative candidate genes influencing the trait. The large QTL region is likely the result of low levels of recombination seen across much of the chromosome which we suspect is due to a large pericentromeric region. Additionally, X chromosomes have less opportunity for recombination in our cross since the X does not cross over in F_1_ males. Our GO enrichment analysis did not reveal any overrepresented terms within the 95% credible interval. However, several of the 211 protein-coding genes in this region have been identified as candidate genes for other relevant phenotypes in *Dendroctonus*. One gene in this region is associated with flight capability in MPB and shows differential expression of a very-long-chain 3-oxoacyl-CoA reductase (LOC109538030) (Shegelski et al. 2021). Another gene, encoding an E3 ubiquitin protein-ligase (LOC109537900), has been implicated in cold tolerance in the closely related species *D. valens* (Zhao et al. 2021). The presence of multiple ecologically relevant genes on the conserved *Dendroctonus* X chromosome suggests this chromosome could play a special role in ecological adaptation in the genus.

Previously, crosses between some MPB populations in the central and western haplogroups with distinct neo-Ys produced sterile hybrid males (Bentz et al., 2011; Bracewell et al. 2011; Bracewell et al. 2017). In this study, we did not observe similar reproductive incompatibilities between the central (AZ) and eastern (UT) haplogroups. Our results demonstrate that some crosses between haplogroups result in sterile hybrid males while others do not. We suspect this is due to variable patterns of neo-Y evolution and degeneration of different haplogroups (Bracewell et al. 2017). However, it is important to note that our cross was only one direction (AZ male x UT female) and the reciprocal was not performed or investigated. Additionally, more sensitive intrinsic sterility phenotypes were not explored, and our results do not rule out extrinsic hybrid sterility which may be particularly relevant given our results and discussed below.

We identified a significant QTL reflecting a long generation time in males hemizygous for the AZ-haplotype (the central haplogroup) and a shorter generation time for the UT-haplotype (the eastern haplogroup). These results corroborate previous common garden experiments using these populations (Bracewell et al., 2013; Soderberg et al., 2021). A recent study using MPB from the central haplogroup in southern Colorado and the eastern haplogroup in northern UT further resolved the physiological traits associated with observed differences in generation time between the haplogroups. Hansen et al. (2024) revealed a diapause in the adult stage of central haplogroup individuals that was not manifest in the eastern haplogroup population. Diapause is a dormancy that can be induced by one or multiple environmental cues, including temperature and photoperiod, that facilitates synchrony of life stages with optimal conditions (Tauber 1986). The intraspecific diapause phenotype can be highly variable, particularly among populations that have adapted to local climates (Pruisscher et al. 2018; Ragland et al. 2019). There is empirical evidence that the adult diapause is mediated by thermal input (Soderberg et al. 2021; Bentz et al. 2022). The diapause phenotype, associated with the AZ population (central haplogroup) appears partially recessive to the non-diapause phenotype at multiple individual markers in our data.

We did not detect a significant QTL for adult body size although previous studies have found a genetic basis for the trait (Bentz et al. 2001; Bracewell et al. 2013). One challenge with our study was that our parent populations had only minor differences in body size (∼ 0.24 mm) and measurement error may have reduced our ability to detect a QTL. It is also likely the underlying genetic architecture of body size in MPB is rather complex and involves many genes of small effect, similar to what has been found in *Drosophila melanogaster* (Carreira et al. 2009). In either case, future work using more distinct MPB populations with increased sample sizes may improve the power of QTL identification.

Our coverage analysis revealed heterogenous patterns of neo-X/neo-Y differentiation, suggesting neo-Y chromosome degeneration in MPB is not uniform. Males appear hemizygous from ∼48 Mb to the end of the chromosome, a region identified as being the conserved ancestral-X using synteny comparisons with other beetles. Previous work also identified this region as the ancestral-X, although estimating the start position at 51.1 Mb (Keeling et al. 2022). Similar to results from Keeling et al., (2022) we found the first ∼25 Mb of the X chromosome shows greater neo-X/neo-Y sequence similarity. The variable patterns we observe for neo-X and neo-Y differentiation suggests the loss of recombination between these young sex chromosomes likely occurred in different stages in the past.

Although one in four described animal species on the planet is a beetle, rates and patterns of recombination (i.e., the recombination landscape) in beetles are not well known. In our study, we identified regions of low recombination and low gene density near the ends of multiple autosomes and in the middle of the X chromosome. These patterns likely reflect underlying chromosomal features, such as pericentromeric/heterochromatic regions. Pericentromeric regions are often enriched for satellite sequences and transposable elements (Sun et al. 1997; Naish et al. 2021). They are also often found to be negatively correlated with recombination rate (Rizzon et al. 2002; Nambiar and Smith 2018) and gene density (Wright et al. 2003). Comparisons of our linkage map with that of Keeling et al. (2022) are largely consistent, suggesting recombination patterns extend to other populations and reflect an underlying feature of the MPB genome.

Future research is needed to identify putative centromere and pericentromere sequence and structure in MPB. Additionally, a broader investigation of genomic patterns across beetles will enhance our understanding of the evolution of genome structure in beetles as a whole.

Our linkage map revealed putative inversions on chromosomes 2, 3, and the X, relative to the reference genome assembly (Keeling et al. 2022). There are two possible explanations for these patterns. First, individuals used to construct our map could be fixed for alternate chromosomal configurations. Alternatively, scaffolding errors could be present in the published reference genome. For the X chromosome, we believe the published orientation is likely incorrect. This region was previously found to have very low levels of recombination (Fig. 4), making scaffolding via linkage challenging, and Hi-C heatmaps actually support an inverted orientation similar to our linkage map (Keeling et al. 2022). For putative inversions found on chromosomes 2 and 3, it is likely these are real. Inversions segregating in MPB have been identified by others, and the populations used to scaffold the reference genome were from a geographically distant population (Keeling et al. 2022). Structural variation is also a common feature of genomes (Buchanan and Scherer 2008) and can be geographically structured and play a role in adaptation (Kirkpatrick and Barton 2006; Akopyan et al. 2022). Intriguingly, recent work has identified numerous segregating inversions in the bark beetle *Ips typographus*, some of which may be associated with diapause (Mykhailenko et al., 2024). The adaptive significance of large scale structural variation is receiving renewed interest (Akopyan et al. 2022) and MPB is well poised to explore the role of these variants in adaptation and speciation.

One of the more surprising results was the difference found in F_2_ male and female linkage maps. The difference was driven entirely by the X chromosome. While previous work also found a larger X chromosome linkage map in females (Keeling et al. 2022), we found that this was largely due to erroneous markers that do not align well to the reference genome. In contrast, our data show the difference spans the entire X chromosome and is not due to misaligned markers. The lack of crossovers found in F_2_ males in the region from 28 - 43 Mb only partially explains the pattern, since the pattern holds on the ancestral-X as well. One possible explanation for our observation is that loci on alternative haplotypes are incompatible and therefore a crossover in this region leads to a combination of alleles that causes male lethality.

Under this scenario, only F_2_ males that lack the recombined haplotype would survive and be included in our genotyping and phenotyping step. However, if this were true, we would also expect to see a female skewed sex ratio given that some subset of males are dying during development. Unfortunately, our rearing methods lead to fairly inaccurate estimates of total brood number and sex ratio, and we actually observed slightly more males than females (132 males and 111 females) which is not consistent with this hypothesis. The observed sex ratio in natural populations is typically female-biased and thought to be driven by differences in thermal tolerance due to body size (Reid 1958; McGhehey 1968; Lachowsky and Reid 2014). It is plausible that our experimental design lacks the power to detect a significant bias in sex ratio, even if it exists. Although our incompatibility hypothesis is highly speculative, it is hard to envision other scenarios where F_2_ males inherit X chromosomes that differ from those of F_2_ females. An additional complication is the presence of the neo-Y, which is rather large and appears to have many functional genes (Bracewell et al. 2017; Dowle et al. 2017; Horianopoulos et al. 2018). All F_2_ males inherited an AZ neo-Y, and previous analyses suggest the UT and AZ neo-Y likely differ in gene content that could lead to incompatibilities with other loci (Bracewell et al. 2017). Future work, such as reciprocal crosses and sampling in natural populations, will be needed to disentangle the complexities of this intriguing pattern and test the male lethality hypothesis.

## DATA AVAILABILITY

Raw sequence reads are available under BioProject ID PRJNA1111460 via NCBI (https://www.ncbi.nlm.nih.gov/). All relevant code is available at https://github.com/cpushman/MPB-QTL-project.

## ACKNOWLEDGEMENTS

We thank Jeff Good for providing guidance on early stages of the QTL analysis. We thank Karen Mock for helping with experimental design. We thank Matt Hansen and Jim Vandygriff for collecting MPB populations and for performing the cross. Jacqueline Lopez and the Notre Dame Genomics & Bioinformatics Core Facility for assistance with library preparation and QC. This work would have been impossible without help from the Indiana University UITS high-performance computing cluster. The findings and conclusions in this report are those of the author(s) and should not be construed to represent any official USDA or U.S. Government determination or policy. Ryan Bracewell was supported by the NIH NIGMS MIRA grant: R35GM151123.

